# The major diagnostic VSG LiTat 1.3 of the human parasite *Trypanosoma brucei gambiense* is a trimer in solution

**DOI:** 10.1101/2025.10.28.684996

**Authors:** Niki Danel, Rob Geens, Israel Mares-Mejía, Wim Versées, Jan Van Den Abbeele, Yann G.-J. Sterckx

## Abstract

Human African trypanosomiasis (HAT) remains a significant health burden in sub-Saharan Africa, with serological diagnosis relying heavily on parasite variant surface glycoproteins (VSGs). In this study, we present evidence that LiTat 1.3 (a key VSG in the diagnosis of *T. b. gambiense* infections) displays a homotrimeric architecture in solution instead of the archetypal homodimeric structure expected for a VSG. This was demonstrated by adopting an integrative structural biology approach encompassing AlphaFold-based structure prediction, analytical gel filtration (AGF), size exclusion chromatography with multi-angle light scattering (SEC-MALS), and small-angle X-ray scattering (SAXS). Furthermore, the SAXS data demonstrate that the C-terminal domains of trimeric VSGs exhibit the same degree of flexibility as observed in dimeric VSGs. Hence, the biophysical characterization of LiTat 1.3 VSG adds to the limited, yet growing body of knowledge that certain VSG classes occur as homotrimers instead of homodimers.

**Author’s Summary:** Human African trypanosomiasis (HAT) is caused by *Trypanosoma brucei gambiense*, a parasite transmitted by tsetse flies. To survive in the human host, these parasites cover themselves with a coat consisting of millions of identical copies of surface proteins called variant surface glycoproteins (VSGs). This VSG coat is regularly switched by the parasite to escape the immune system. Some of these VSGs, including one known as LiTat 1.3, are used in diagnostic tests to detect potentially infected patients. In our study, we discovered that, unlike most VSGs that form pairs of identical molecules (homodimers), LiTat 1.3 assembles into groups of three (homotrimers). Using structural and biophysical techniques, we showed that this trimeric form is stable in solution and retains the dynamic behavior observed in dimeric VSGs. Understanding how such structural variations arise and how they influence immune recognition may help explain why certain VSGs, like LiTat 1.3, are particularly effective in diagnosis and could ultimately guide the development of improved tools to monitor and control sleeping sickness.

## Introduction

Human and animal African trypanosomiasis (HAT and AAT, respectively) are neglected tropical diseases caused by protozoan parasites of the *Trypanosoma* genus. The *T. brucei* species harbors three subspecies: *T. b. gambiense* and *T. b. rhodesiense* (both human-infective and responsible for HAT, alias sleeping sickness) and *T. b. brucei* (non-infective to humans and cause of AAT, alias nagana). Since transmission solely relies on blood-feeding tsetse flies (*Glossina* spp.), the geographical range of these parasites is correlated with the spread of the insect vector, which is typically confined to foci in sub-Saharan Africa. While *T. b. rhodesiense* causes acute infections, *T. b. gambiense* (gHAT) cases are chronic and responsible for 94% of reported HAT cases (2024). Infection by either subspecies leads to fatal neurological symptoms if left untreated [1,2].

African trypanosomes are extracellular parasites that can survive in their vertebrate hosts for months or even years, despite being continuously exposed to the immune system. Once metacyclic trypomastigotes are inoculated into the vertebrate host through the bite of an infected tsetse fly, the parasites transform into bloodstream forms (BSFs) giving rise to a systemic infection. The BSFs thrive within the host blood circulation but they also colonize the lymphatics, skin, brain, testes, adipose tissue, and lungs [3–7]. Being extracellular parasites, BSFs rely on sophisticated mechanisms to evade host immune responses. Among these, the antigenic variation is the most notorious in which trypanosomes employ a controlled mechanism of switching their major surface antigens called variant surface glycoproteins (VSGs) [8–12]. Although trypanosomes have a repertoire of >1000 VSG-encoding genes, individual parasites only express a single gene at a time, thereby resulting in the display of identical VSGs at their surface. On the latter, the expressed VSGs form a dense monolayer of around 10^7^ identical molecules that acts as a protective barrier to shield underlying invariant proteins from immune recognition. Each VSG coat is called a variant antigen type (VAT), and trypanosomes expressing the same VSG belong to the same VAT. VSGs are highly immunogenic and readily elicit an effective humoral response, in which antibodies matured against the exposed VAT lead to parasite clearance. Antigenic variation entails that trypanosomes switch their VSG coat by successively cycling their *VSG* gene repertoire, either by activating a different intact *VSG* gene or by generating novel mosaic VSGs. This leads to the emergence of parasite subpopulations that escape recognition by existing antibodies, giving rise to new parasitemic waves [12,13] The structure-function relationship of VSGs has received much attention given their importance in trypanosome immunobiology. Despite a low amino acid sequence identity, all VSGs possess a conserved structural architecture: i) an N-terminal domain (NTD; 300-400 residues) consisting of a membrane-distal top lobe separated from a membrane-proximal bottom lobe by a 3-helix stalk, and ii) a C-terminal domain (CTD; 80-120 residues) harboring a glycosylphosphatidylinositol (GPI) anchor, which attaches the VSG to the parasite surface [14]. VSGs are further categorized into two classes (A and B) based on the arrangement of the bottom lobe in their primary sequences. Class A includes all VSGs studied to date that form homodimers, with their bottom lobe residues positioned at the C-terminal end of the NTD. In contrast, class B comprises VSGs that form concentration-dependent homotrimers, with their bottom lobe residues located at the N-terminal end. Furthermore, class B VSGs frequently feature *O*-linked carbohydrate modifications, a trait that has not yet been observed in class A VSGs [14–16]. Hence, three features of VSG structure underpin antigenic variation: i) the NTDs display poor sequence conservation (∼10-30% sequence identity) and adopt a myriad of distinct topologies, ii) VSGs may undergo post-translational modifications (e.g., *O*-linked glycosylation) with potent immunomodulatory effects [14–16], challenging the belief that antigenic variation is solely driven by amino acid sequence divergence, and iii) VSGs may adopt distinct oligomeric states (thereby challenging the longstanding paradigm that VSGs solely occur as homodimers). Studies on the solution structure of homodimeric VSGs have revealed that, while the NTDs behave as rigid entities, the CTDs are hallmarked by a high degree of flexibility, thereby rendering the entire VSG coat highly dynamic [17,18]. While the same behavior could be expected for homotrimeric VSGs, such evidence is yet to be presented.

As the clinical signs of gHAT are not specific enough for an unambiguous diagnosis, sensitive and specific diagnosis plays a key role in identifying true positives. Since the 1970s, serological tests relying on the detection of VAT-specific antibodies have formed the cornerstone of gHAT diagnosis and are key for the successful reduction in disease incidence during the last decades. These serological gHAT diagnostic tests all rely on antibody reactivity towards two predominant *T. b. gambiense* VSGs: the Lille Trypanozoon antigen types 1.3 and 1.5 (LiTat 1.3 and LiTat 1.5, respectively) [19–21]. Given their extensive diagnostic application for gHAT screening/diagnosis, an in-depth knowledge of the solution state structure of these key *T. b. gambiense* VSGs represents an added value. While LiTat 1.5 was recently characterized as an archetypical dimeric class A1 VSG [18], the structural details of LiTat 1.3 remain poorly understood. Here, we present the application of an integrative structural biology approach to investigate the solution state of *T. b. gambiense* LiTat 1.3 VSG. By combining AlphaFold structural modeling, analytical gel filtration (AGF), size exclusion chromatography coupled to multi-angle light scattering (SEC-MALS) and small-angle X-ray scattering (SAXS), our findings provide compelling evidence that LiTat 1.3 exists as a homotrimer in solution instead of a homodimer. Furthermore, the SAXS-based modeling shows that the LiTat 1.3 solution behavior is best described by a conformational ensemble due to a highly flexible CTD, thereby indicating that the CTDs found in trimeric VSGs exhibit the same degree of structural plasticity as those present in VSG dimers. Our results therefore suggest that, besides serving as a means to expand the concept of antigenic variation, trimeric VSGs also contribute to the parasite’s highly dynamic surface antigen coat.

## Materials and methods

### Ethics statement

Experimental protocols for the cultivation of *T. b. gambiense* were reviewed and approved by the Ethical Committee of the Institute of Tropical Medicine, Antwerp, Belgium (approval codes TTP-2021- 1 and TTP-2021-2).

### Protein production and purification

The purified VSG LiTat 1.3 was provided by the Applied Technology and Production unit at ITM and was part of a large VSG batch that was used for the production of the gHAT-RDT diagnostic test (HAT Sero K-SeT) by Coris BioConcept (Gembloux). To initiate the VSG production, *T. b. gambiense* bloodstream forms expressing LiTat 1.3 were cultivated in rats after intraperitoneal injection of 10^8^ trypanosomes/mL, administered at 1 mL per 100 g body weight. Approximately 72 hours after initiating the rat infection, when parasites are in their first expanding growth phase, blood was collected and passed over a self-packed diethylaminoethyl (DEAE)-52 column (8 L bed volume) to retain the trypanosomes. Following sample loading, the column was washed with phosphate-saline-glucose (PSG; 83 mM D-(+)-Glucose, 38 mM Na_2_HPO_4_, 29 mM NaCl, 2 mM NaH_2_PO_4_, pH 8.0) until the eluate was clear. Elution was carried out at a fixed flow rate (maintained at approximately one drop per second) using a peristaltic pump and the collected trypanosome-containing fraction was kept on ice. Subsequently, the trypanosome suspension was concentrated and subjected to two washing steps in PSG via sequential centrifugation (1,000 x g and 4°C). The concentrated parasite suspension was aliquoted and centrifuged again. Following the removal of the supernatant, the resulting 1 mL pellets were stored at -80°C.

For VSG extraction and purification, the trypanosome pellets were resuspended in five volumes of ice- cold Concanavalin A (ConA) binding buffer (500 mM NaCl, 1 mM MgCl₂ 1 µM CaCl₂, 10% (v/v) phosphate buffer (100 mM Na₂HPO₄ titrated with 100 mM NaH₂PO₄ to pH 8.0) and transferred to 10 mL centrifuge tubes on ice. The suspension was vortexed to ensure homogeneity, incubated on ice for 15 min, and centrifuged (42,000 x g for 2 h at 4°C). The supernatant was transferred to a new tube and the pellet was resuspended in 3 mL of ice-cold ConA binding buffer, vortexed and centrifuged again. The resulting supernatants were combined, filtered (0.22 µm filter), and stored at 4°C. The next day, the VSG-containing supernatant was loaded onto a 1 mL HiTRAP ConA 4B column (Cytiva) at a flow rate of 0.5 mL/min, after which the column was washed with 10 column volumes (CVs) of ConA binding buffer, and the VSG was eluted using ConA elution buffer (ConA binding buffer supplemented with 10% methyl α-D-mannopyranoside). Then, the VSG eluate was concentrated to approximately 10 mg/mL and subsequently buffer switched to either HEPES-buffered saline (HBS) or phosphate-buffered saline (PBS), using a 30-kDa cutoff centrifugal filter (Centricon) and stored at -80°C.

### Analytical gel filtration

Analytical gel filtration (AGF) was performed using a Superdex 200 Increase GL 10/300 column (Cytiva) coupled to an ÄKTA FPLC system. The column was pre-equilibrated with HBS (25 mM HEPES, 150 mM NaCl, pH 7.4). Samples of 100 µL were injected and eluted at a flow rate of 0.75 mL/min. The Bio-Rad gel filtration standard was run under the same conditions for molecular mass estimation. Relevant elution fractions were collected for SDS-PAGE analysis and stored at -80°C.

### SDS-PAGE

Purified VSG samples were mixed with 4x SDS sample buffer and heated at 100°C for 10 min to ensure complete denaturation. Samples were loaded on a 4-15% mini-PROTEAN TGX precast gel (Bio-Rad) and electrophoresis was performed in Tris-Glycine-SDS buffer (Bio-Rad) at 120 V. Protein bands were visualized using PageBlue protein staining solution (Thermo Scientific) with the Precision Plus Protein Dual Color standard (Bio-Rad) allowing molecular mass estimations.

### Small-angle X-ray scattering and ensemble modelling

Small-angle X-ray scattering (SAXS) was conducted at the BioSAXS beamline SWING (SOLEIL, Gif-sur-Yvette, France) [22]. SEC-SAXS data were acquired on a Shodex KW404-4F column, pre-equilibrated in 20 mM HEPES, 150 mM NaCl, 3% glycerol, pH 7.4. A sample of 50 µL (15.51 mg/mL) was injected and subsequently eluted at a flow rate of 0.2 mL/min. Scattering data were collected with an exposure time of 750 msec and a dead time of 10 msec, calibrated to absolute units using the scattering of pure water [23]. Data processing and analysis was performed using the ATSAS package and BioXTAS RAW [24–26]. The information on data collection and derived structural parameters is summarized in Supplementary Table 1.

Molecular models of LiTat 1.3 dimers and trimers were generated using AlphaFold-Multimer[27] based on the primary amino acid sequence (GenBank accession no. KJ499460.1) [28]. The signal peptide, which is cleaved from the mature VSG protein, was predicted using the SignalP 6.0 tool [29] and excluded from subsequent modeling. Theoretical scattering curves of the AlphaFold2 models and their respective fits to the experimental data were calculated using FoXS [30]. SAXS-based ensemble modelling was carried out using BilboMD [31,32]. For all runs, 800 conformations were generated per R_g_ bin, with minimal and maximal R_g_ values set at 7% and 35% of the experimentally determined R_g_, respectively. The overall goodness-of-fit between the final models and the experimental data are reported through the calculation of a χ2 value, with N_k_ being the number of points, σ(q_j_) the standard deviations, and c a scaling factor.

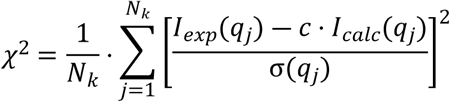

Furthermore, comparisons of theoretical and experimental scattering curves include the presentation of a residuals plot (Δ/σ vs. q, where Δ indicates the difference between experimental and calculated intensities), which enables a local inspection of the model fit to the data.

Visualization and structural analysis of molecular models were conducted using UCSF ChimeraX [33].

### Size exclusion chromatography with multi-angle light scattering

Purified VSG LiTat 1.3 was analyzed by size exclusion chromatography using an Agilent Bio SEC-3 column coupled to an HPLC Alliance system (Waters) equipped with a 2998 PDA detector (Waters), a

TREOS II MALS detector (Wyatt technology) and a RI-501 refractive index detector (Shodex). The samples were prepared at two concentrations (0.5 mg/mL and 5 mg/mL) in PBS. A volume of 50 µL of each was injected onto a pre-equilibrated column in the same buffer at a constant flow rate of 0.2 mL/min. The Astra 7.3.0 software (Wiatt technology) was used to analyze the data, and a BSA sample (1 mg/mL) was used to normalize and align the signals of the different detectors, using a dn/dc value of 0.185 mg/g.

## Results

### *In silico* structure predictions of LiTat 1.3 VSG favor a homotrimeric arrangement

The LiTat 1.3 NTD was first classified using a Hidden Markov Model (HMM)-based Python pipeline[34], which automatically assigns the most probable subtype (A1–A3, B1–B2) based on the highest-scoring sequence alignment against HMM profiles derived from previously typed VSGs. This HMM-based analysis assigned LiTat 1.3 to the B2 subtype, which harbors trimeric VSGs [14]. Next, we employed AlphaFold-Multimer to predict structural models for both dimeric and trimeric configurations of LiTat 1.3. Consistent with the assignment of LiTat 1.3 as a B type VSG, a comparative analysis reveals that the trimeric structural model is favored, as it is supported by superior local and global validation metrics (**Fig. 1A**). Especially the NTDs and their interaction interfaces are predicted with great confidence as advocated by the predicted aligned error (PAE) plots and AlphaFold-Multimer confidence scores. Interestingly, the CTDs are characterized by significantly lower pLDDT scores, which implies that these may be intrinsically disordered [35]. Finally, a structural superposition of the LiTat

**Figure 1.**
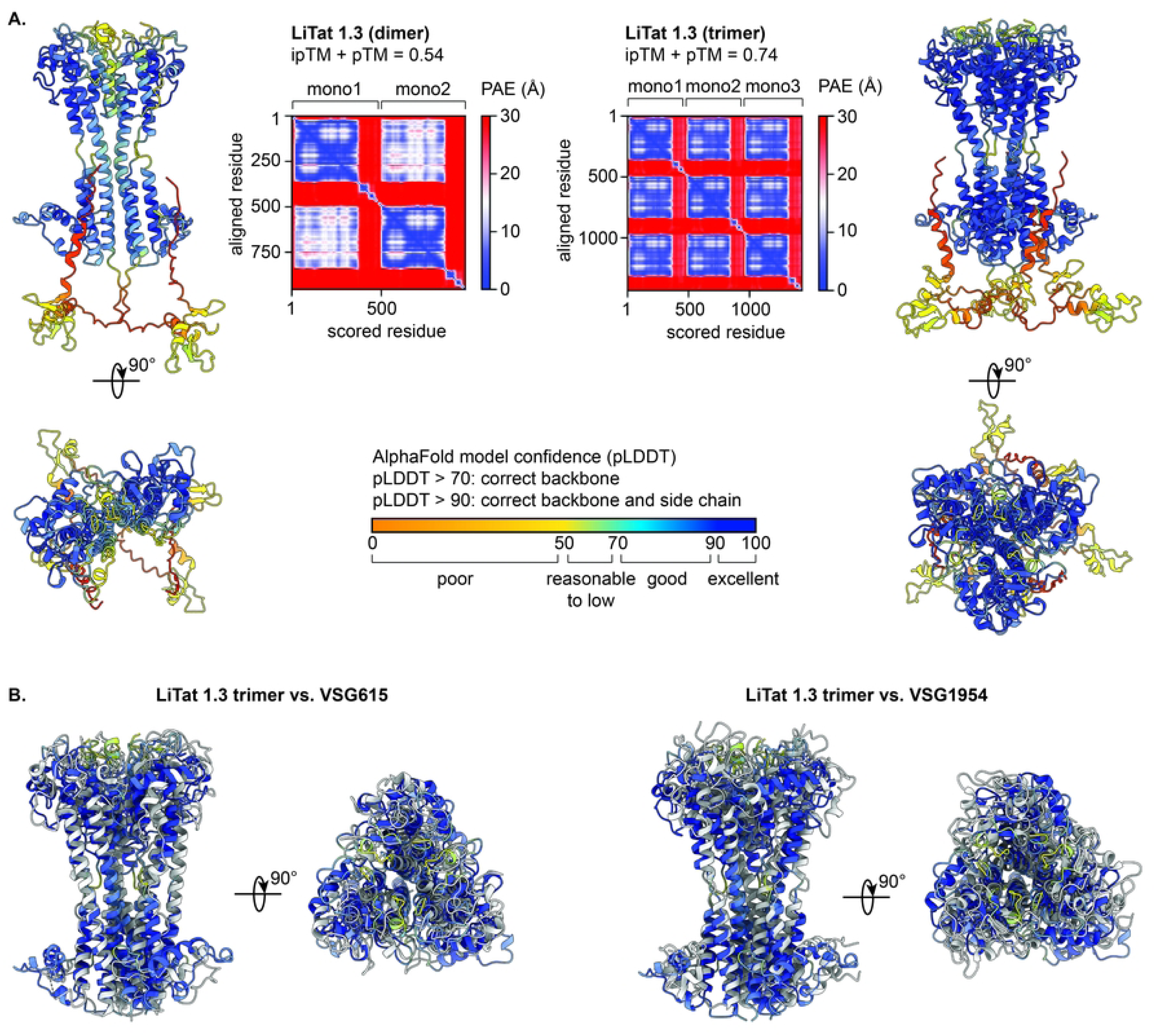
AlphaFold-based structure prediction of LiTat 1.3 and analysis of the obtained models. (A.) Cartoon representations of the LiTat 1.3 dimer (left) and trimer (right) models generated by AlphaFold. The models are colored according to the predicted local distance difference test (pLDDT) score, which reflects (local) model quality as indicated by the legend at the bottom. For both models, the predicted aligned error (PAE) and AlphaFold-Multimer model confidence (0.8*ipTM + 0.2*pTM) are shown. The PAE provides a distance error for every residue pair, and is calculated for each residue x (scored residue) when the predicted and true structures are aligned on residue y (aligned residue). The pTM (score between 0 and 1) provides a measure of similarity between two protein structures (in this case, the predicted and unknown true structure) over all residues and thus reports on the accuracy of prediction within a single chain. The interface pTM (ipTM, score between 0 and 1) provides a measure of similarity between two protein structures (in this case, the predicted and unknown true structure) over only interfacing residues and thus reports on the accuracy of prediction for a complex. (B.) Structural overlay of the NTDs of i) the AlphaFold LiTat 1.3 trimer model and the experimentally determined structure for VSG615 (left), and ii) the AlphaFold LiTat 1.3 trimer model and the experimentally determined structure for VSG1954 (right). In both cases, the AlphaFold LiTat 1.3 model is colored according to the pLDDT score as in panel A, while the experimentally determined structures are colored in light gray.

1.3 trimer AlphaFold model with crystal structures of other Class B2 VSGs [14,16] indeed shows a similar structural arrangement at the NTD trimer interfaces (Figure 1B; Cα RMSD values of 2.26 Å and 3.66 Å for respectively). Hence, the *in silico* analysis favors a homotrimeric arrangement for LiTat 1.3 VSG.

### LiTat 1.3 VSG occurs as a homotrimer in solution

Next, we employed various in solution biophysical techniques to validate the *in silico* predictions: analytical gel filtration (AGF), SEC coupled to multi-angle light scattering (SEC-MALS), and small-angle X-ray scattering (SAXS).

AGF was performed with LiTat 1.3 VSG at different concentrations (Figure 2A). Analysis of the chromatograms consistently reveals the existence of two elution peaks with apparent molecular masses (MM_app_) of ∼280 kDa and ∼100 kDa, respectively. Both elution peaks contain LiTat 1.3 VSG as evidenced by SDS-PAGE analysis (Fig. 2A, inset). Interestingly, the first elution peak represents the major elution fraction at all tested concentrations, indicating that LiTat 1.3 VSG most probably adopts a homo-oligomeric architecture in solution. As AGF determines MM_app_ based on hydrodynamic volume, it is inherently sensitive to protein shape and conformational flexibility in solution. Hence, extended and/or highly dynamic conformations lead to MM_app_ estimations that are higher than would be expected based on sequence alone, making it difficult to draw any conclusions on the exact oligomeric state of LiTat 1.3 VSG under the two observed elution peaks. To further investigate the oligomeric state of LiTat 1.3 VSG in solution, we performed SEC-MALS under the same conditions (Figure 2B). SEC-MALS is a technique that enables MM determination on an absolute scale, as it is not influenced by protein shape and/or conformational flexibility. The elution profiles measured at two concentrations show elution peaks with average absolute MM values close to those theoretically expected for a homotrimer, thereby providing strong evidence that LiTat 1.3 VSG occurs as a homotrimer in solution. Interestingly, in both AGF and SEC-MALS experiments, the elution profiles remain unaltered regardless of the tested concentration, suggesting that LiTat 1.3 homotrimer formation is concentration-independent within the tested concentration range.

**Figure 2.**
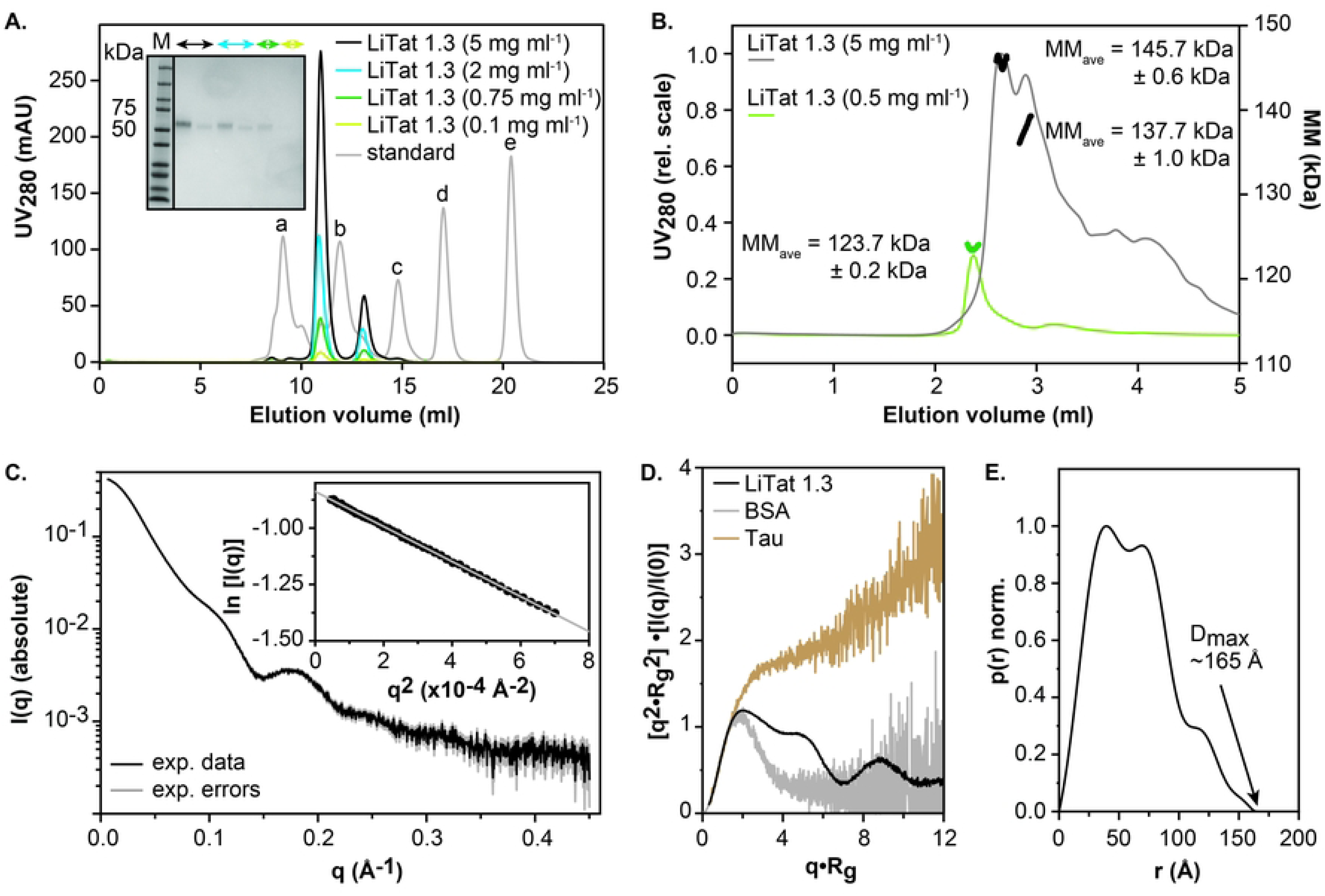
In-solution biophysical characterization of LiTat 1.3. (A.) AGF. The elution profiles of LiTat 1.3 injected at various concentrations onto a Superdex 200 Increase GL 10/300 column are shown in light green (0.1 mg/mL), green (0.75 mg/mL), blue (2 mg/mL), and black (5 mg/mL). SDS-PAGE analyses of the fractions under the collected peaks are shown in the inset. The elution profile of the standard is indicated by the grey line, with elution peaks indicated by small letters: a) bovine thyroglobulin (MM = 670 kDa), b) bovine γ-globulin (MM = 158 kDa), c) chicken ovalbumin (MM = 44 kDa), d) horse myoglobulin (MM = 17 kDa), e) vitamin B12 (MM = 1.35 kDa). (B.) SEC-MALS. The elution profiles (plotted on a relative scale on the first y-axis) of LiTat 1.3 injected at various concentrations onto an Agilent Bio SEC-3 column are shown in green (0.5 mg/mL) and grey (5 mg/mL). The molecular masses determined for the corresponding elution peaks (plotted on an absolute scale on the second Y-axis) are indicated by the green and black dots, respectively. The average molecular masses and their standard deviations are shown for the reader’s convenience. (C.) SAXS. The experimental data and errors are shown as black and grey lines, respectively. The inset displays the Guinier region with the experimental data and the Guinier fit visualized as black dots and a grey line, respectively. (D.) Normalized Kratky plot. The data of LiTat 1.3 VSG, BSA (reference globular protein), and Tau (reference random-coil IDP) are colored black, grey, and brown, respectively. (E.) Pair-distance distribution function. The maximal particle dimension (D_max_) is shown for the reader’s convenience.

Finally, the in-solution features of LiTat 1.3 VSG were also probed by SAXS (Figure 2C), which is a (low- resolution) structural technique that is highly suitable for studying the conformational behavior of biological macromolecules in solution [36,37]. A prerequisite for the extraction of structural parameters from a SAXS data set is that the sample under study is ideal (there are no interparticle effects) and mono-disperse (the sample is homogeneous), which can be assessed through Guinier analysis. The experimentally collected LiTat 1.3 VSG scattering curve obtained from the first (major) elution peak is characterized by a long, linear Guinier region, which is strongly indicative of ideal, monodisperse behavior (Figure 2C, inset). A thorough analysis of SAXS-derived structural parameters and MM estimations (Supplementary Table S1) are fully consistent with a homotrimer, thereby directly supporting the AGF and SEC-MALS data sets. Interestingly, the SAXS data indicate that the LiTat 1.3 VSG homotrimer is characterized by a significant degree of flexibility. First, normalized Kratky plot analysis (Figure 2D)[38] reveals an in-solution behavior that is intermediate compared to reference curves for random coil intrinsically disordered proteins (IDPs; Tau) and rigid, globular proteins (bovine serum albumin, BSA). While the latter have a bell-shaped appearance with a tail returning to zero, those for random coil IDPs reach a plateau followed by a continuous increase with higher scattering angles. The normalized Kratky plot of LiTat 1.3 displays multiple peaks (consistent with the occurrence of multiple domains and an oligomeric state) and contains a tail that does not return to zero (indicating a certain degree of flexibility). Second, the p(r) plot also suggests a pronounced asymmetry reflected by an elongation ratio larger than 2 (Figure 2E)[39].

In conclusion, these in-solution biophysical studies provide experimental evidence that LiTat 1.3 VSG adopts a homotrimeric, non-globular architecture in solution.

### The solution structure of LiTat1.3 VSG is best described by a conformational ensemble that considers CTD flexibility

Next, the SAXS data were employed to investigate the conformational dynamics of LiTat 1.3 VSG in solution. A first comparison between the experimental scattering curve and a scattering curve calculated for the initial AlphaFold-Multimer single-state model clearly demonstrates that the in-solution behavior of LiTat 1.3 VSG cannot be explained by a single conformer (χ² = 167.87; **Figure 3**). Together with the above-mentioned biophysical data suggesting that LiTat 1.3 VSG contains a certain degree of flexibility, this indicates that its solution structure is best described by a conformational ensemble. To this end, we relied on SAXS-driven ensemble modeling with BilboMD as previously performed for dimeric VSGs [18]. Here, we assumed that the CTD would be highly flexible / intrinsically disordered given that i) this was suggested by AlphaFold-Multimer modeling (*cfr*. Figure 1) and ii) this behavior has been thoroughly documented for dimeric VSGs [17,18]. Indeed, ensemble modeling that considers a highly flexible CTD provides a significantly better fit (χ² = 2.03), thereby suggesting that LiTat 1.3 adopts a conformational ensemble rather than a single, rigid structure. From the obtained models, it becomes clear that the CTD undergoes substantial structural rearrangements in solution and possesses a conformational freedom akin to the CTDs of dimeric VSGs. For the sake of completion, we calculated scattering curves for both rigid and flexible LiTat 1.3 VSG dimers and compared these to the experimental data set. It again becomes clear that LiTat 1.3 does not occur as a dimer in solution and that the homotrimeric arrangement best explains the data set.

**Figure 3.**
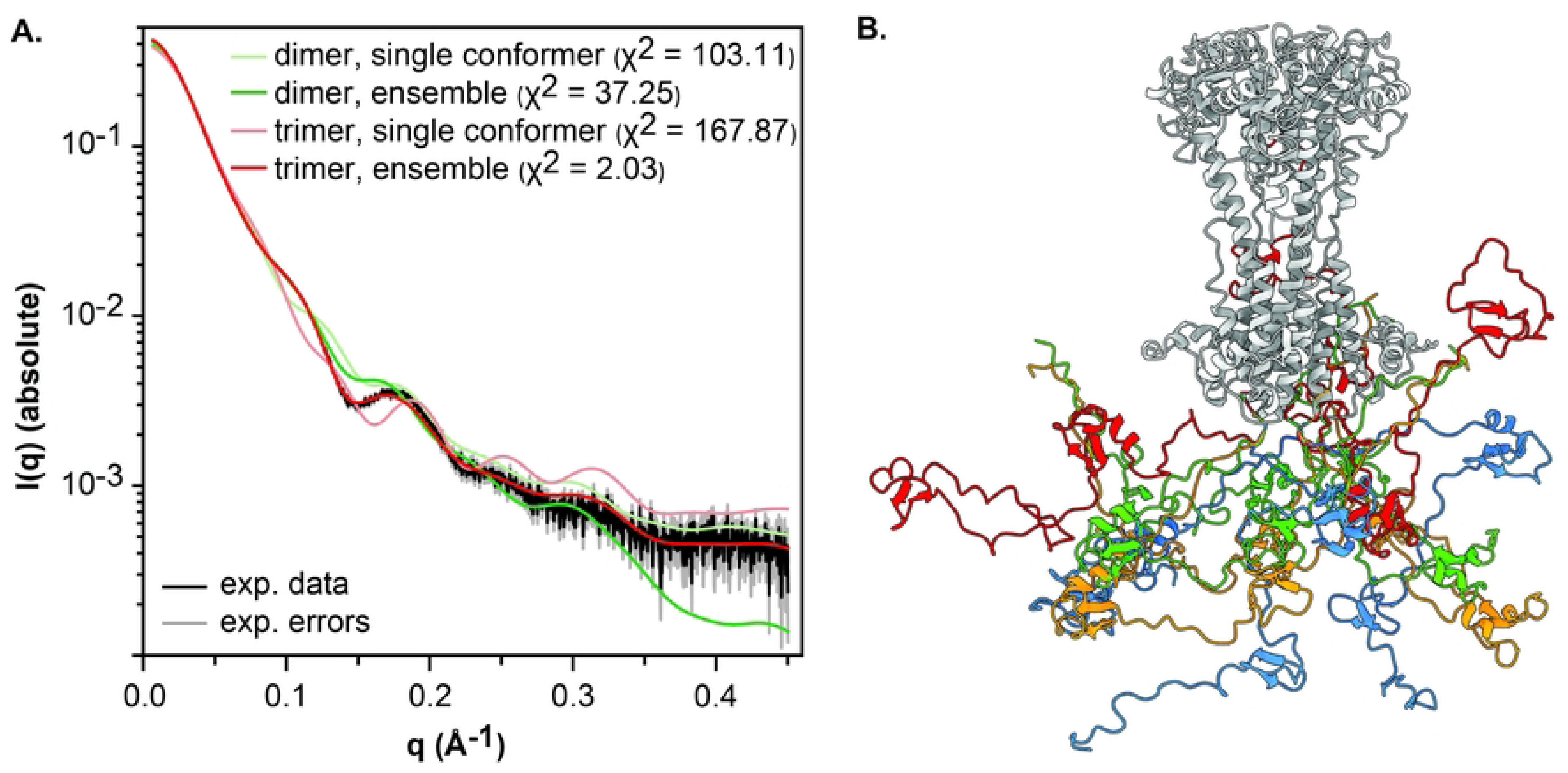
Conformational ensemble of the LiTat 1.3 trimer. (A.) Comparison of the fits for different models to the experimental data. The latter are visualized in black, with the error margins shown in grey as in Figure 2. The fits of the different models are shown in light green (LiTat 1.3 dimer, AlphaFold single conformer), green (LiTat 1.3 dimer, conformational ensemble), salmon (LiTat 1.3 trimer, AlphaFold single conformer), and red (LiTat 1.3 trimer, conformational ensemble). The χ^2^ are reported as a measure for the goodness of fit for the reader’s convenience. (B.) Cartoon representation of the LiTat 1.3 trimer conformational ensemble that best describes the in-solution SAXS data. All conformers have been superposed using the NTD, which is colored grey. The CTD of each conformer is shown in a different color to highlight the CTD’s high flexibility.

## Discussion

*T. b. gambiense* is the primary causative agent of HAT, a parasitic disease that continues to impose a significant public health burden in sub-Saharan Africa. Accurate diagnosis remains critical for controlling transmission and advancing disease elimination efforts [40]. In this context, the *T. b. gambiense*-specific LiTat 1.3 VSG is a key antigen for serological diagnosis. Indeed, LiTat 1.3 expressing parasites are the sole basis for the Card Agglutination Trypanosomiasis Test (CATT)[41] that is used for massive screening of populations at risk, and it is the major *gambiense*-specific antigen in the individual HAT Sero K-Set rapid diagnostic test (Coris BioConcept)[42]. Despite its critical role in serological gHAT- diagnostics, the structural and biophysical properties of LiTat 1.3 have remained largely unexplored. In this study, we comprehensively characterized its structural features, oligomeric state, and conformational flexibility.

Our data demonstrate that LiTat 1.3 VSG purified from *in vivo* grown trypanosome BSFs exists as a homotrimer in solution, as evidenced by a combination of biophysical analyses (AGF, SEC-MALS, and SAXS) and structural modeling (SAXS-based conformational modeling using AlphaFold-Multimer models as starting structures). Interestingly, under the tested experimental conditions, LiTat 1.3 appears to maintain its trimeric state, which is in contrast with previously characterized class B2 trimeric VSG-species, where oligomerization was found to be concentration-dependent [14,43]. These results suggest that the LiTat 1.3 trimer may exhibit an enhanced stability compared to these other trimeric VSGs, although the underlying mechanism for this stability remains unclear. Furthermore, the SAXS data and SAXS-based modeling demonstrate that in-solution behavior of LiTat 1.3 is best described by a conformational ensemble rather than a rigid, singular structure. The ensemble contains both compact and extended conformers that exhibit structural features that mirror those observed in dimeric VSGs; *i.e.*, a rigid, structured NTD and a highly flexible CTD of which its dynamic character is mediated by intrinsic disorder [17,18]. Although CTD conformational flexibility was so far only reported for dimeric VSGs, our findings provide the first experimental evidence that this feature is also present in trimeric VSGs. This demonstrates that CTD dynamics are not limited to dimeric forms and may represent a more general property of the VSG family, regardless of their oligomeric state. Hence, as this observed flexibility is most likely also present on the BSFs surface membrane, it indicates that trimeric VSGs also contribute to the dynamics of the VSG coat [17,18].

Previous studies have shown that there is a strong bias towards a presentation of class B (homotrimeric) VSG on the parasite surface in natural *T. b. gambiense* infections, a characteristic not observed in other trypanosome species [44] This raises the question of why *T. b. gambiense* favors such a VSG coat configuration. This preference may be linked to specific structural and functional benefits offered by homotrimeric VSGs to sustain chronical infections in the human host (as is observed for *T. b. gambiense*), although this remains speculative and enigmatic.

In summary, our findings provide new insights into the structural and biophysical properties of LiTat 1.3, the predominant diagnostic antigen for *T. b. gambiense*. We demonstrate that LiTat 1.3 forms a stable, concentration-independent trimeric structure, classifying it to a class B2 VSG. Future high-resolution structural studies, including those that explore the VSG in complex with antibodies, will be essential for further identifying the major epitopes that are involved in immune recognition of the LiTat 1.3 VSG and to unravel the underlying basis of this VSG as the key diagnostic antigen for gHAT. Here, comparative analyses with other *T. b. gambiense* VSGs may also help clarify why LiTat 1.3 dominates current serodiagnostics. However, the full explanation for its diagnostic predominance likely involves a combination of structural, genomic, and evolutionary factors that merit further investigation. In conclusion, this work contributes to the broader understanding of VSG structural diversity, particularly in terms of oligomerization and domain flexibility, and may help frame future investigations into whether these properties play a role in antigenicity or diagnostic performance.

## Acknowledgements

The authors wish to thank the staff of the SWING beam line at SOLEIL Synchrotron (Aurélien THUREAU) for the outstanding beam line support. We thank Ana Lucia Fajardo Castañeda and Nicolas Bebronne from the Applied Technology and Production unit at ITM, led by Caroline Rombouts, for generously providing the purified LiTat 1.3 VSG used in this study. The authors acknowledge the use of the CalcUA and VSC supercomputing facilities and wish to thank the staff for their outstanding support. N.D. was supported by a University of Antwerp BOF “Seal of Excellence” (code 53274) and is a doctoral fellow supported by the FWO-Vlaanderen (1143426N). The research was supported by a Strategic Research Program Financing from the Vrije Universiteit Brussel (SRP95 to W.V.) and the Bill and Melinda Gates Foundation (code INV031353) as part of the improvement of gambiense-HAT diagnostics.

## Supporting information

**Table S1.** SAXS data collection and scattering-derived parameters for LiTat 1.3 VSG.

|  | *T. brucei gambiense* LiTat 1.3 VSG |
| --- | --- |
| **Data collection parameters** |  |
| Beam line | SWING (SOLEIL) |
| Wavelength (Å) | 0.99 |
| $q$ range (Å<sup>-1</sup>)* | 0.00045 – 0.55 |
| Concentration (mg ml<sup>-1</sup>) (mode) | 15.5 (SEC-SAXS) |
| Buffer conditions | 20 mM HEPES, 150 mM NaCl, 3% glycerol, pH 7.4 |
| Temperature (°C) | 20 |
| **Structural parameters<sup>§</sup>** |  |
| $I(0)$ (cm<sup>-1</sup>) [from Guinier] | 0.43 |
| $R_g$ (Å) [from Guinier] | 48.21 |
| $I(0)$ (cm<sup>-1</sup>) [from $p(r)$] | 0.44 |
| $R_g$ (Å) [from $p(r)$] | 46.64 |
| E.R. | 2.76 |
| $D_{max}$ (Å) | ~165 |
| Porod volume estimate, $V_p$ (Å<sup>3</sup>) | 252354 |
| Porod exponent | 3.8 |
| **Molecular mass determination** |  |
| MM (kDa) [from SAXSMoW on final merged curve] | 147.00 ($q = 0.25 \text{ \AA}^{-1}$) |
| MM (kDa) [from $Q_R$ on final merged curve] | 142.25 ($q = 0.25 \text{ \AA}^{-1}$) |
| MM (kDa) [from $V_p/1.7$] | 148.44 |
| Calculated MM from sequence (kDa) | 144.50 |
| **Ensemble modeling** |  |
| Conformer generation and selection | BILBOMD |
| Protein regions selected to be fixed/rigid | 24-366, 384-417, 434-457 |
| **Software employed** |  |
| Data processing and analysis | ATSAS, BioXTAS RAW |
| Computation of theoretical intensities and fitting | FoXS, MultiFoXS |
| **SASBDB entry** | SASDXA9 |

Abbreviations: $I(0)$ , extrapolated scattering intensity at zero angle; $R_g$ , radius of gyration calculated using either Guinier approximation (from Guinier) or the indirect Fourier transform package GNOM [from $p(r)$ ]; $MM$ , molecular mass; $D_{max}$ , maximal particle dimension; $V_p$ , Porod volume; $V_{ex}$ , particle excluded volume; E.R., elongation ratio \*Momentum transfer $|q| = 4\pi\sin(\theta)/\lambda$

